# Interpretable machine learning with tree-based shapley additive explanations: application to metabolomics datasets for binary classification

**DOI:** 10.1101/2022.09.19.508550

**Authors:** Olatomiwa O. Bifarin

**Author notes:** School of Chemistry and Biochemistry, Georgia Institute of Technology, Atlanta GA, United States of America.

## Abstract

Machine learning (ML) models are used in clinical metabolomics studies most notably for biomarker discoveries, to identify metabolites that discriminate between a case and control group. To improve understanding of the underlying biomedical problem and to bolster confidence in these discoveries, model interpretability is germane. In metabolomics, partial least square discriminant analysis (PLS-DA) and its variants are widely used, partly due to the model’s interpretability with the Variable Influence in Projection (VIP) scores, a global interpretable method. Herein, Tree-based Shapley Additive explanations (SHAP), an interpretable ML method grounded in game theory, was used to explain ML models with local explanation properties. In this study, ML experiments (binary classification) were conducted for three published metabolomics datasets using PLS-DA, random forests, gradient boosting, and extreme gradient boosting (XGBoost). Using one of the datasets, PLS-DA model was explained using VIP scores, while a tree-based model was interpreted using Tree SHAP. The results show that SHAP has a more explanation depth than PLS-DA’s VIP, making it a powerful method for rationalizing machine learning predictions from metabolomics studies.

## 1 Introduction

Metabolomics aims to report a global snapshot of a biological system’s metabolic status (1). The adequate tools to answer this question give rise to large datasets inherently. Therefore, the need to engage in classical and modern statistics, with machine learning being an example of the latter. Classical statistics aim to draw inferences about the population from a sample, while machine learning (ML) finds a generalizable predictive pattern in a dataset without explicit instructions (2, 3). ML is used in experimental metabolomics workflows to find predictive patterns in data, for example, for biomarker discoveries (4, 5). However, the interpretation of these predictive models has mainly been limited to the use of partial least squares discriminant analysis (PLS-DA) in metabolomics (6, 7).

PLS-DA, also known as a projection on latent structures, combines features from principal component analysis (PCA) and multiple linear regression (8). It extracts latent variables, the best predictors, from the independent variables and project results to a lower-dimensional space, as in PCA. PLS has been the standard multivariate analysis algorithm used in metabolomics for two main reasons. One, given the structure of the metabolomics dataset, a large number of features *vs.* smaller sample sizes, latent variables’ projection onto a smaller dimensional space allows for its utility as a feature selection algorithm. Two, the linear regression structure inherent with the algorithm makes for an interpretability method, the variable importance in projection (VIP) scores (9). As such, PLS-DA is an interpretable ML algorithm that models the linear latent covariance with the feature’s matrix (***X***) and the response matrix (***Y***). Despite the popularity of PLS-DA in metabolomics, the linear relationship assumption and the solely global interpretability it affords are obvious limitations.

Some of the best-performing machine learning methods are notoriously black box models (10, 11) i.e., models that are not interpretable. Linear models like linear regression and PLS-DA are interpretable because of their linear assumptions. In general, intrinsically interpretable models are so because of their simple structures, lending themselves to features such as the weights in linear models and the learned tree structure in decision trees. However, biological data can have non-linear relationships (12), which might require models to learn more complex relationships in such datasets for better performance. One approach to explain complex, black-box models is to apply interpretation methods after machine learning modeling. These methods are called post-hoc interpretable machine learning (IML) methods, and they can be model agnostic (13).

Additionally, despite the interpretability property of intrinsically interpretable models, they are usually limited to global explanations – explaining the entire model behavior, rather than local explanations that explain individual predictions. In general, a ML interpretable method with local and global interpretations (high representativeness) is indeed desirable. Other desirable properties of explanation methods include high expressive power (the *language* of expression), low algorithmic complexity, and high fidelity (the accuracy of the interpretation) (14). There are several IML method like partial dependence plot (PDP) (15), individual conditional expectation (ICE) (16), accumulated local effects (ALE) (17), permutation feature importance (18, 19), and local interpretable model-agnostic explanations (LIME) (20). However, only SHapley Additive exPlanations (SHAP) gives a solution that satisfies the quality of high representativeness, fidelity, and high expressive power (21, 22). Thus, it was the method of choice in the study. The method has been empirically verified (23), and it has been applied to many fields of study, including medicine (23, 24), cheminformatics (25, 26), and ecology (27).

In this article, 1) Shapley values and SHAP for tree-based models (TreeSHAP) were introduced, 2) the classification performance of PLS-DA and tree-based models (random forest, gradient boosting, and XGBoost) were compared using three clinical metabolomics data sets, and finally, 3) one of the tree ensemble model was explained with the aid of Tree SHAP (22). In brief, Tree SHAP aided both local and global explanations.

## 2 Results

### 2.1 Shapley Additive Explanations

#### 2.1.1 Shapley Values

In a standard metabolomics experimental workflow (28), after metabolic measurements and appropriate spectral and data processing, the resultant data matrix can be used for modeling using machine learning (Fig 1a). Shapley Additive exPlanations (SHAP) allows for the local interpretations of predictions by calculating each metabolomic feature importance score for each sample prediction. Additionally, SHAP is used to derive an accurate global interpretation of the model, giving rise to its high representativeness as a post-hoc IML method (Fig 1b). SHAP is based primarily on Shapley values, a cooperative game theory method (21). Developed by Lloyd Shapley, it is a fair and axiomatically unique method of attributing rewards from a cooperative game (29). Where a *game* is a machine learning model, each metabolomic feature values are *players* in a *game,* and the predicted class membership of the sample is the *outcome* of the *game;* Shapley value gives a unique solution to fairly attribute the contributions of each *player* to the outcome of the *game*. The Shapley value defines the feature importance of feature value *i* in the equation below and Fig 1c:

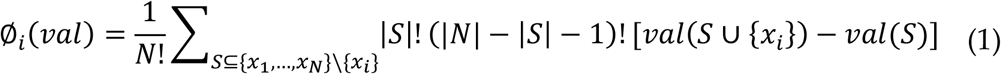

**Fig 1.**
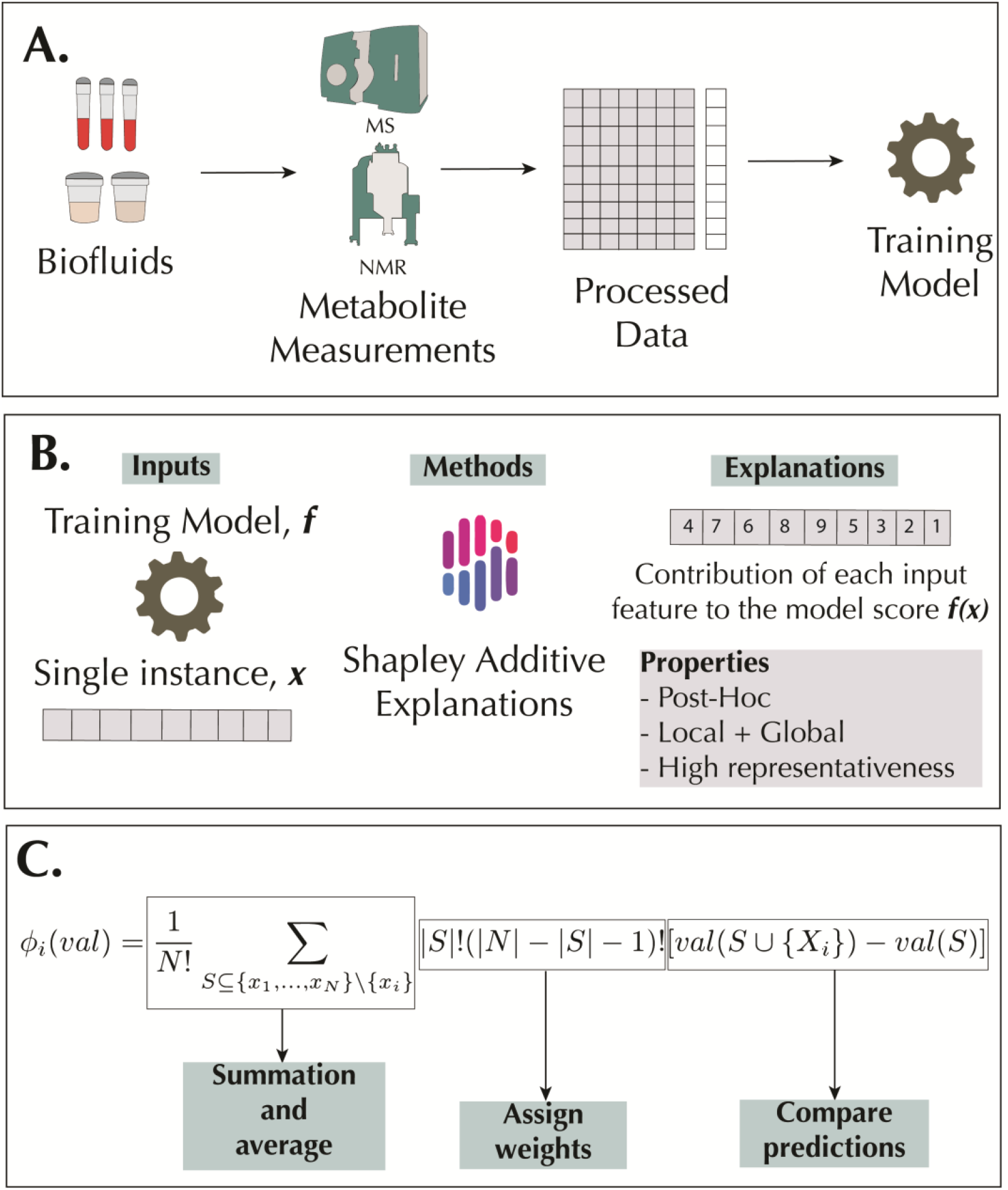
Metabolomics workflow and SHAP methodology. A: A metabolomics workflow that culminates with model training for predictive or regression purposes. B: SHAP allows for local and global interpretations of model predictions. Explanations are made locally, and because of the additivity property of Shapley values, the methods allow for global interpretations. C: A sample calculation of Shapley values of a feature *x_i_*.

Where *S* is a subset of features in the model, *val*(*S*) corresponds to the model output for *S, N* is the total number of features, and *x* is the feature values for the sample to be explained, that is *x* ≜ {*x*_1_…,*x_N_*} ∈ ℝ^*N*^. In brief, the marginal contribution of *x_i_* is given by *val*(*S* ∪ {*x_i_*}) – *val*(*S*). Weights are assigned to these marginal contributions by the different ways the sub-set could be formed before the addition of *x_i_*: |*S*|! and after the addition of *x_i_*: (|*N*| – |*S*| – 1)!. The summation over all the possible sets *S* is conducted, followed by the average 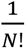.

The Shapley value is a unique solution because it satisfies the axioms of symmetry (or consistency), dummy (or null effect), and additivity (or local accuracy) (29). Symmetry implies that if the marginal contribution of the metabolomic feature values *x_z_* and *x_k_* is the same, the Shapley value attributed to each feature value will also be the same.

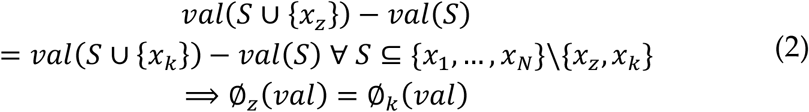

Dummy implies that if a feature value *x_z_* do not impact a model, the Shapley value attributed will be zero.

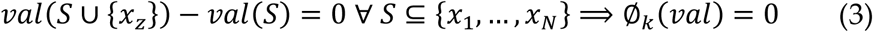

Finally, local accuracy means that the summation of the Shapley values of all feature values in the model equals the model output. As such, the total contributions of all feature values will equal the impact of all feature values on the model output minus the impact with no feature value, mathematically expressed below:

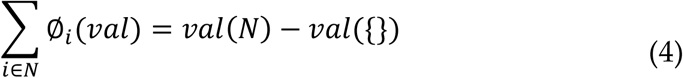

#### 2.1.2 Tree SHAP

The choice of tree-based SHAP ML interpretations for this study was based on the relatively low computational complexity of computing Shapley values and the exact Shapley value that result from the computation (22). However, other methods for approximately Shapley values for non-tree-based methods exist (21), making it a model agnostic IML method. Two problems emanate from the computation of naïve Shapley values, namely, 1) handling of missing features when computing marginal contributions and 2) exponential computational time and algorithmic complexity. First, when computing Shapley values for a feature value, the absence of the feature must be defined while computing the marginal contributions. This is not straightforward in machine learning as opposed to game theory. The problem has been addressed in kernel SHAP (21) by simulating missing features *via* random sampling by replacing the missing value with a fixed value. This invariably creates a sampling-based estimation variance problem; on the other hand, kernel SHAP can be applied to any ML model. In Tree SHAP, exact Shapley values are calculated by ignoring the decision paths of the missing features in the decision tree, getting rid of sampling-based estimation variance in the process. Second, because Tree SHAP computes Shapley values by keeping track of tree transversals to prevent repetition (22), it gives rise to an efficient algorithm by reducing algorithmic complexity from exponential time (*O*(*TL*2^*N*^)) to polynomial time (*O*(*TLD*^2^)). Where *T* is the number of trees, *L* is the maximum number of leaves in any tree, *N* is the number of features, and *D* is the maximum tree depth.

### 2.2 Machine Learning Pipeline and Performance

ML models were built for three metabolomics datasets, as summarized in Table 1 (See methods for details). All tasks were binary classification problems, and since the primary goal of the study is to show the utility of SHAP, a simplified machine learning workflow was used (Fig 2).

**Table 1.** Metabolomics datasets used in the study.

| Dataset | MTBLS404 | MTBLS547 | ST000369 |
| --- | --- | --- | --- |
| **Analytical platform** | LC-MS | LC-MS | GC-MS |
| **Sample Type** | Urine | Caecal | Serum |
| **Sample Size** | 184 | 97 | 80 |
| **Subject of classification** | Sex | High-fat diet | Adenocarcinoma |
| **Classes (Size)** | Male/Female<br>(101/83) | Case/Control<br>(46/51) | Case/Control<br>(49/31) |
| **Metabolomic feature number** | 184 | 42 | 69 |
| **Study** | Thevenot et al. 2015 (30) | Zheng et al. 2017 (31) | Fahrman et al. 2015 (32) |

**Fig 2.**
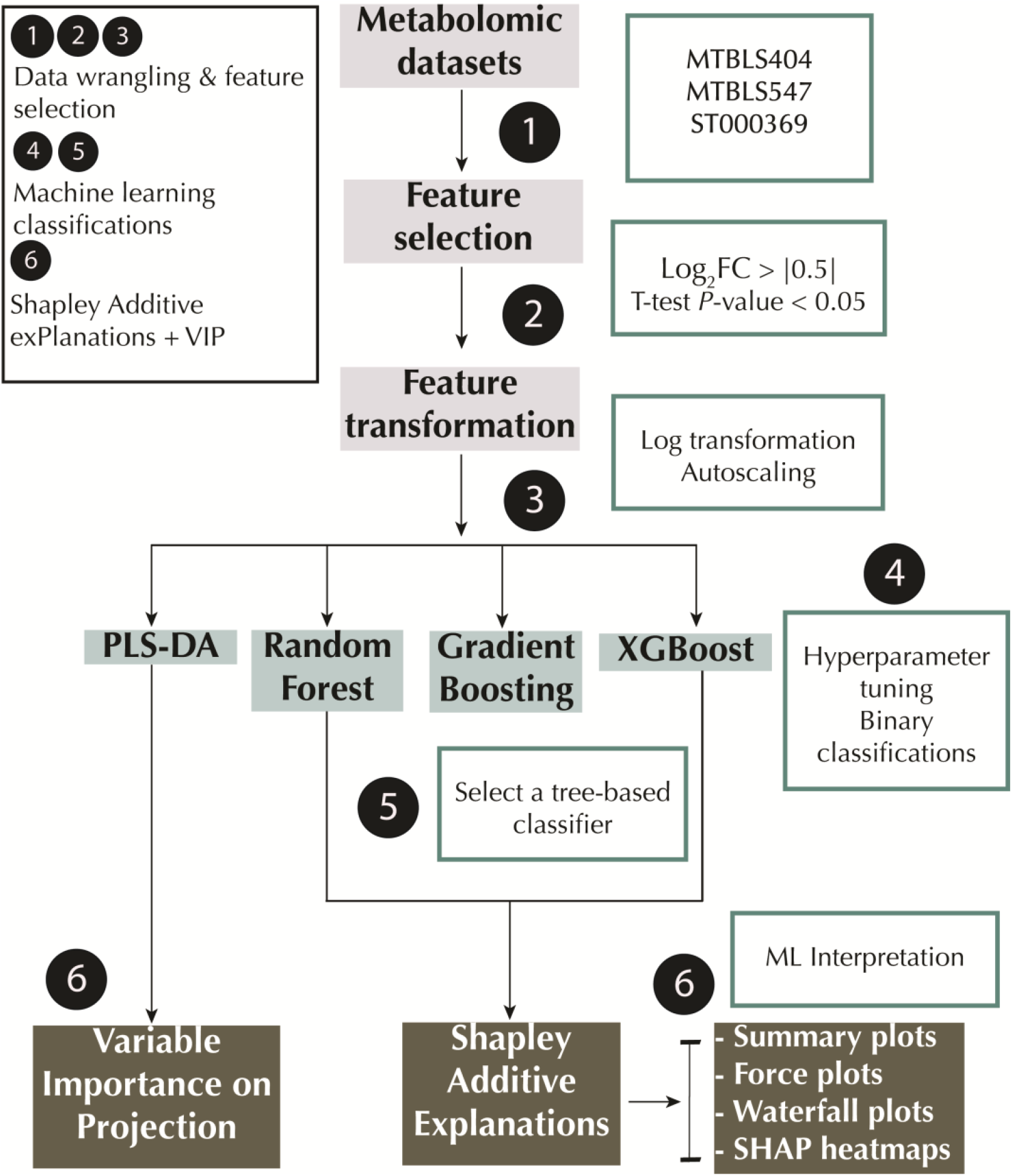
Machine learning pipeline. PLS-DA: Partial Least Square Discriminant Analysis; XGBoost: Extreme Gradient Boosting; VIP: Variable Importance in Projection.

Samples were split into train and test set with 5/6 and 1/6 of the sample size respectively in each set. The train set was used for feature selection and training, while the test set was kept solely for test purposes. Metabolomic features with fold changes greater than 0.5 (Log_2_FC > |0.5|) and *P*-values less than 0.05 (Student’s *T*-test) were selected. Thirty-two features were selected for MTBLS404, 17 features for MTBLS547, and 10 features for ST000369 (S1 Fig). In preparation for the classification tasks, the train and test set were transformed *via* a log transformation followed by autoscaling. Afterward, the selected features using the train sets were used to build machine learning models with PLS-DA, random forest, gradient boosting and XGBoost. Appropriate hyperparameters were tuned (See methods and S1-S3 Tables), and a tree-based classifier was used for Shapley additive explanations. The machine learning performance results of both the baseline and tuned ML method are presented in Table 2. Random forests gave the best predictive scores on MTBLS404 and MTBLS547 test sets with an AUC value of 0.83 and 0.88 respectively. XGBoost, gradient boosting, and PLS-DA models performed as well as random forests on the MTBLS404 dataset with an AUC of 0.83. In addition, an XGBoost model performed as well as random forests on the MTBLS547 dataset with an AUC of 0.88. Finally, a PLS-DA model had the highest AUC value of 0.74 on the ST000369 dataset. Going forward, the MTBLS404 test set will be used to illustrate the utility of SHAP in this study. The dataset is from a urine metabolomics study that investigated human adult urine for variations in age, BMI, and sex (30). In this analysis, the dataset has been used for sex classifications.

**Table 2.**
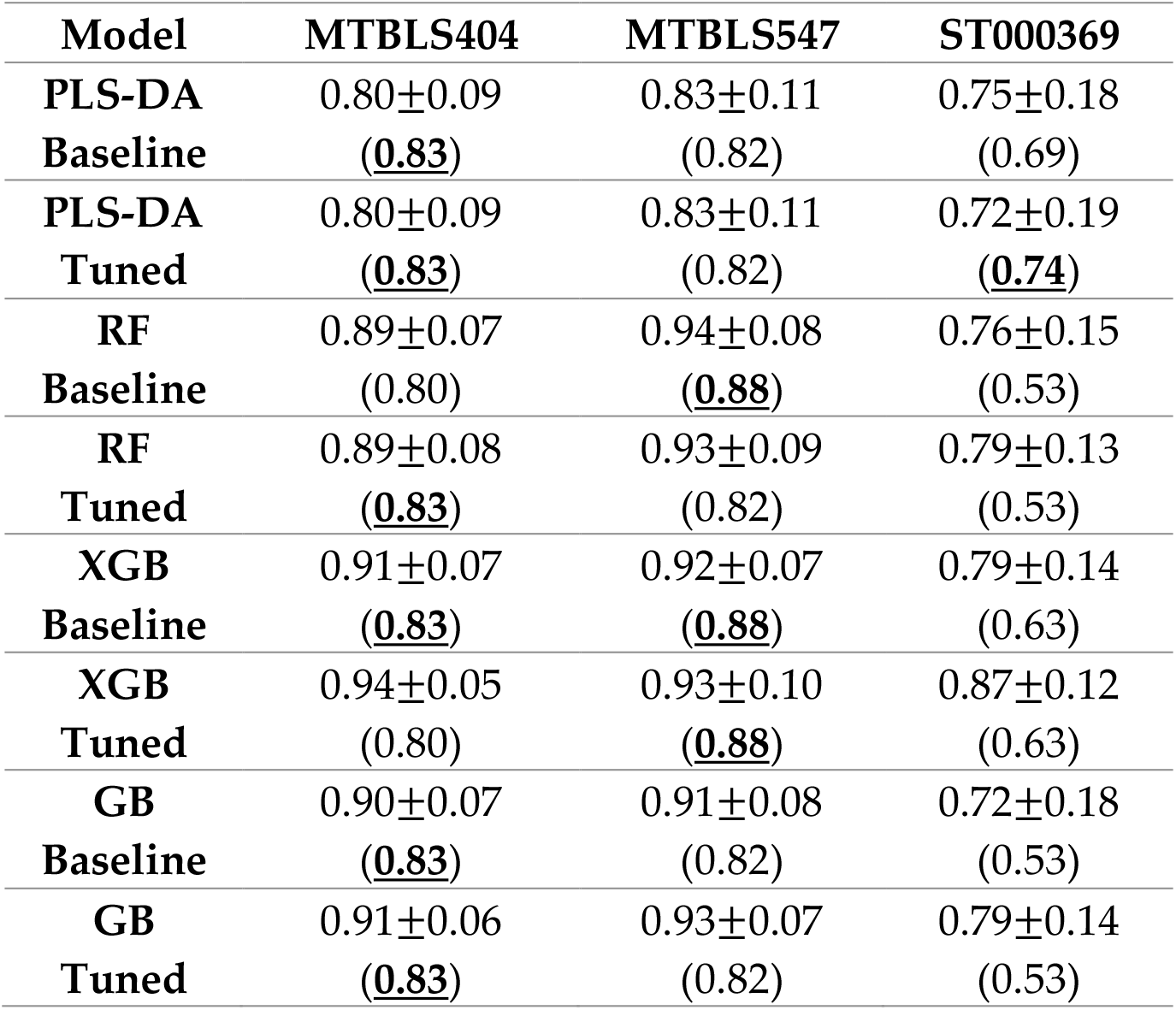
Machine learning performance.

Baseline models use the default hyperparameters in the sci-kit learn library for the python programming language. The tuned models underwent hyperparameter tuning as described in the materials and methods. Predictive scores are the Area Under the Receiver Operating Characteristic Curve (ROC AUC). The training score reports the mean±standard deviation of ROC AUC under 10-fold cross-validation conditions. The test set performance scores are reported in brackets below the train set scores, and the highest test set scores for each dataset are shown in bold texts and underlined. PLS-DA: Partial Least Squares-Discriminant Analysis; RF: Random Forests; XGB: Extreme Gradient Boosting; GB: Gradient Boosting.

### 2.3 Model Interpretations with SHAP

SHAP explanations were computed for the test set of the MTBLS404 dataset trained using random forest with an AUC of 0.83, and these explanations are presented in this section. The PLS-DA VIP score plot, a global interpretation of the PLS-DA model, is shown in Fig 3a. Likewise, Fig 3b shows the SHAP bar plot, displaying the mean of the absolute SHAP values (mean[|SHAP value|]), that is, the average impact of the feature on the model output magnitude. Testosterone glucuronide and *p*-anisic acid have the same feature importance rank on both plots. In addition, the Pearson correlation coefficient of the VIP score and the mean[|SHAP value|] is 0.50 (*p* = 0.03) (Fig 3c), indicating some explanation similarities. On the other hand, the Pearson correlation coefficient of the feature importance attribute of random forests, Gini importance, and mean[|SHAP value|] is 0.99 (*p* = 6.44×10^-25^) (Fig 3d).

**Fig 3.**
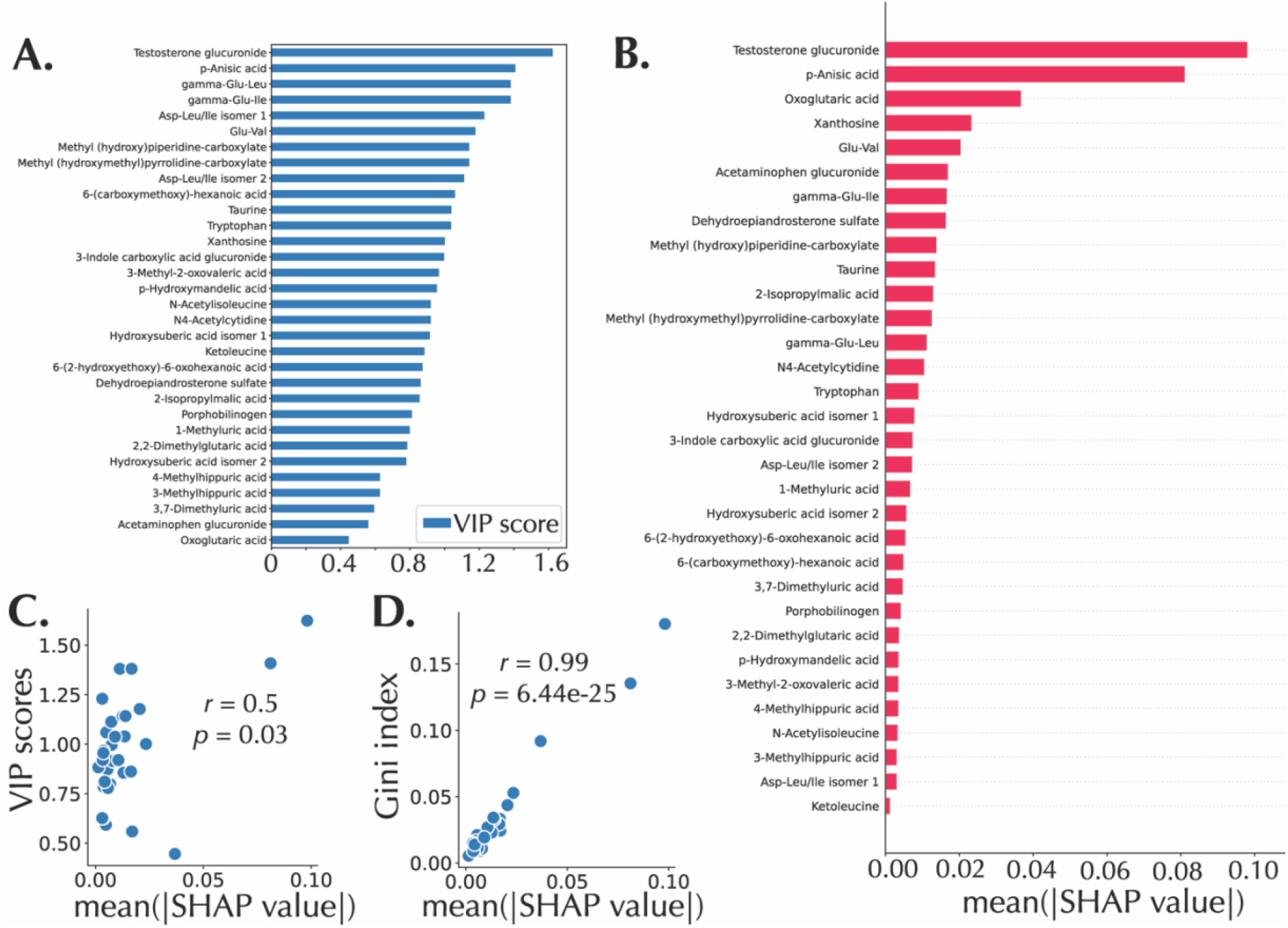
Global feature importance and feature importance correlations. A: PLS-DA VIP score plot. B: SHAP bar plot. C: Scatterplot of the VIP score and the mean(|SHAP value|) with a Pearson’s correlation coefficient of 0.50. D: Scatterplot of the Gini importance score and the mean(|SHAP value|) with a Pearson’s correlation coefficient of 0.99.

In addition, because of the local representativeness property of SHAP, it can give both the global importance score and an explanation of individual predictions in the SHAP summary plot (also called the beeswarm summary plot), enabling a richer visual summarization, as shown in Fig 4a. In the plot, metabolomic features are arranged in a descending order based on relative importance 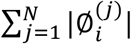, where 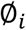 is the Shapley value of feature *i, j* is a sample, and *N* is the total number of samples. Each dot in the summary plot represents a sample plotted against its impact on the model output 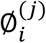. The color of each sample represents the relative abundance of the metabolites, ranging from low (blue) to high (red). Fig 4b displays the most important metabolite in the panel – testosterone glucuronide. High feature values of the metabolite tend to have positive SHAP values, which drives the model to predict males; on the other hand, low feature values of testosterone glucuronide tend to have negative SHAP values, which drives the model to predict the female sex. *p*-anisic acid has an opposite trend, as also shown in Fig 4b. Furthermore, SHAP values was projected in a 2-dimensional space via PCA, which can be used to visualize the impact of a metabolomic feature as observed in both the intensity of the SHAP values and the clustering of positive and negative SHAP values respectively. Fig 4c shows the SHAP embedding plot of testosterone glucuronide with clustering of positive and negative SHAP values, while Fig 4d shows the embedding plot of the least important feature, ketoleucine.

**Fig 4.**
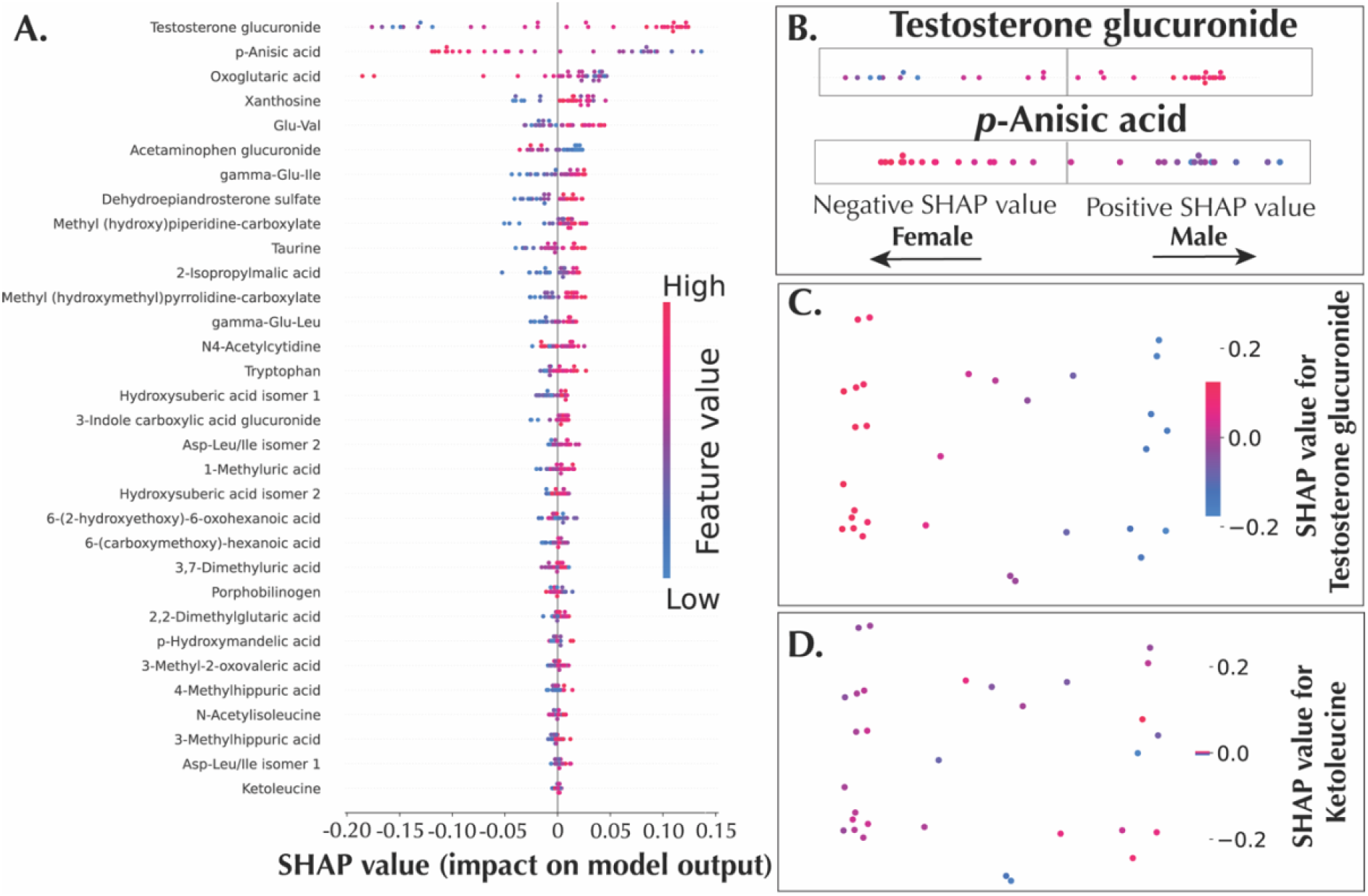
SHAP summary and PCA embedding plots. A: SHAP summary plot. B: SHAP summary plot illustration with testosterone glucuronide with *p*-Anisic acid. C: Embeddings plot highlighting testosterone glucuronide. D: Embeddings plot highlighting Ketoleucine.

Unsupervised clustering techniques are widely used in metabolomics studies to identify groups of classes that cluster together. Because such analyses rely on raw data, which are only processed by a simple standardization technique, it is impossible to cluster samples based on a prediction outcome. On the other hand, SHAP values can be used to generate supervised clustering, where samples are clustered based on explanation similarities for the same prediction outcome. For example, in the heatmap shown in Fig 5, the hierarchical supervised clustering analysis clustered samples that share the same sex prediction for similar reasons, that is, similar metabolomic feature explanations. In this study, the heatmap shows that, more often than not, male sex predictions are associated with positive SHAP values of testosterone glucuronide, and female predictions associate with negative SHAP values of testosterone glucuronide, as evident in the prediction outcome (*f*(*x*)) line plot at the top of the heatmap. This trend indicates that testosterone glucuronide is not only the most important feature in the model. It is also the primary driver of sex predictions, making it a probable candidate for a sole marker of sex, in this dataset.

**Fig 5.**
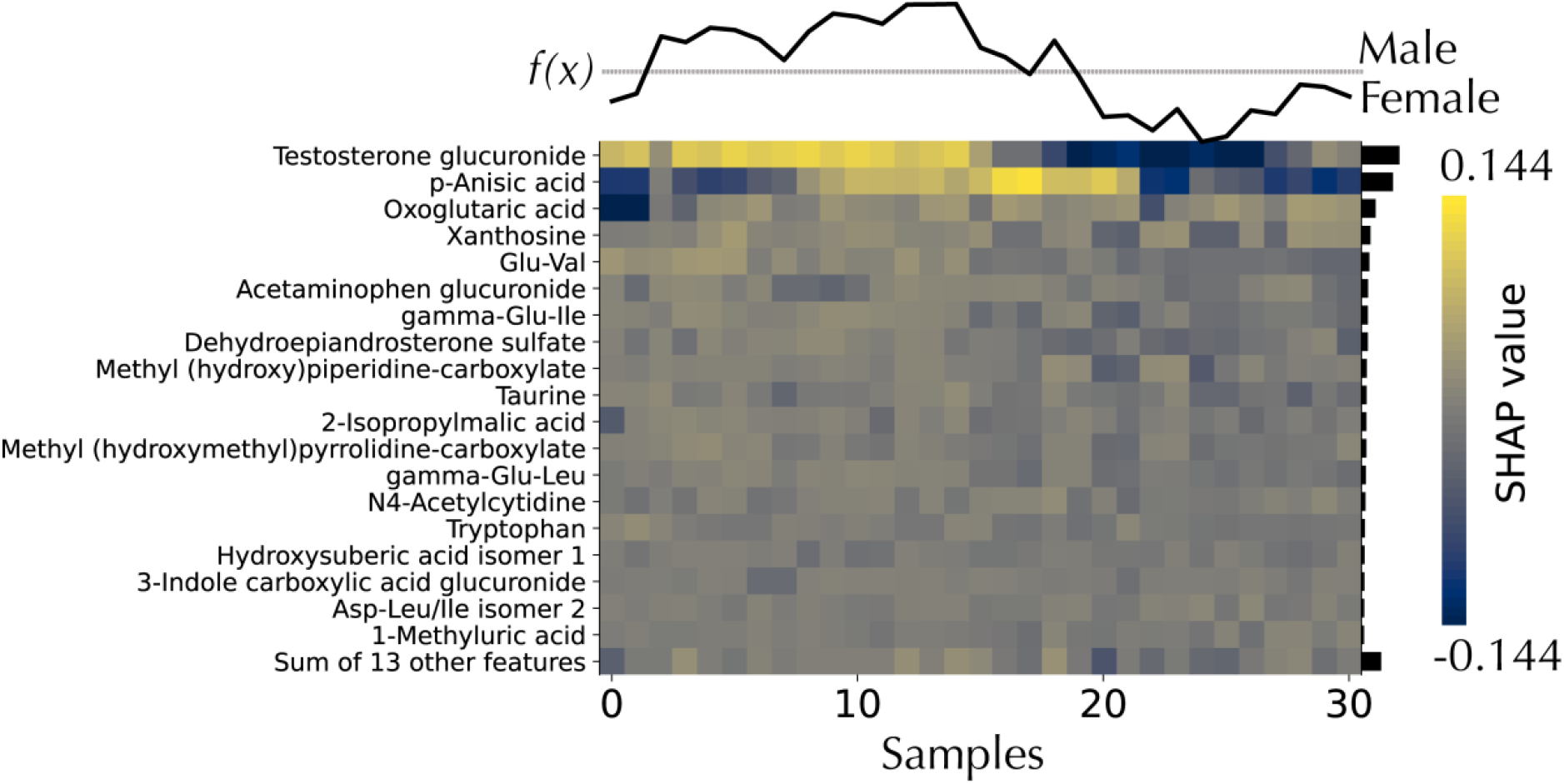
Supervised interpretable hierarchical clustering. Samples are displayed on the x-axis, while features are arranged in ascending order of importance on the y-axis. *f*(*x*) indicates the prediction outcome, with the line plot over the dotted line indicating male predictions, while the line plot below the dotted line indicates female prediction. The bar plot represents the mean absolute SHAP value, the average impact on the model output.

Just like the partial dependence plot (PDP) shows the marginal effect that one or two features have on a model output with a line plot (15), SHAP values can be used to create a better alternative graph called the SHAP dependence plot. The plot shows how the relative abundance of a metabolite changes with the respective impact on the model output (S2 Fig). In addition, dots representing the samples can be colored by the relative abundance of another metabolite to capture the interaction effect if it exists (S2b Fig). Fig S2a displays the relationship between testosterone glucuronide and the SHAP values for the metabolite (the impact of the metabolite on model output), while S2b Fig adds *γ*-glu-leu to the plot by coloring the samples by the relative abundance of *γ*-glu-leu. The S-like curve shows that higher feature values of testosterone glucuronide and *γ*-glu-leu, tend to lead to a male prediction.

Finally, SHAP provides many avenues for local explanations (individual sample explanations), such as the waterfall plot and the force plot visualizers. An example of waterfall plot and force plot are shown in Figs 6a and 6b, respectively. In the waterfall plot (Fig 6b), the x-axis represents the probability of a sample being classified as male, while the y-axis shows the metabolomic features and the respective feature values for the sample. Waterfall plots begin with the expected value of the model output on the x-axis (*E*[*f*(*X*)] = 0.55). This ‘base’ value, 0.55, is the average prediction probability over the test set. The plot displays the impact of the metabolomic features on the model output. The combination of the positive contributions (in red) and the negative contributions (in blue) moves the expected value output to the final model output (*f*(*x*) = 0.80). Positive SHAP values increase the probability of the sample being classified as male, while negative SHAP values decrease the probability of the sample being classified as male. Similarly, the force plot helps visualize each feature’s SHAP values for a given sample with an additive force layout (Fig 6a).

**Fig 6.**
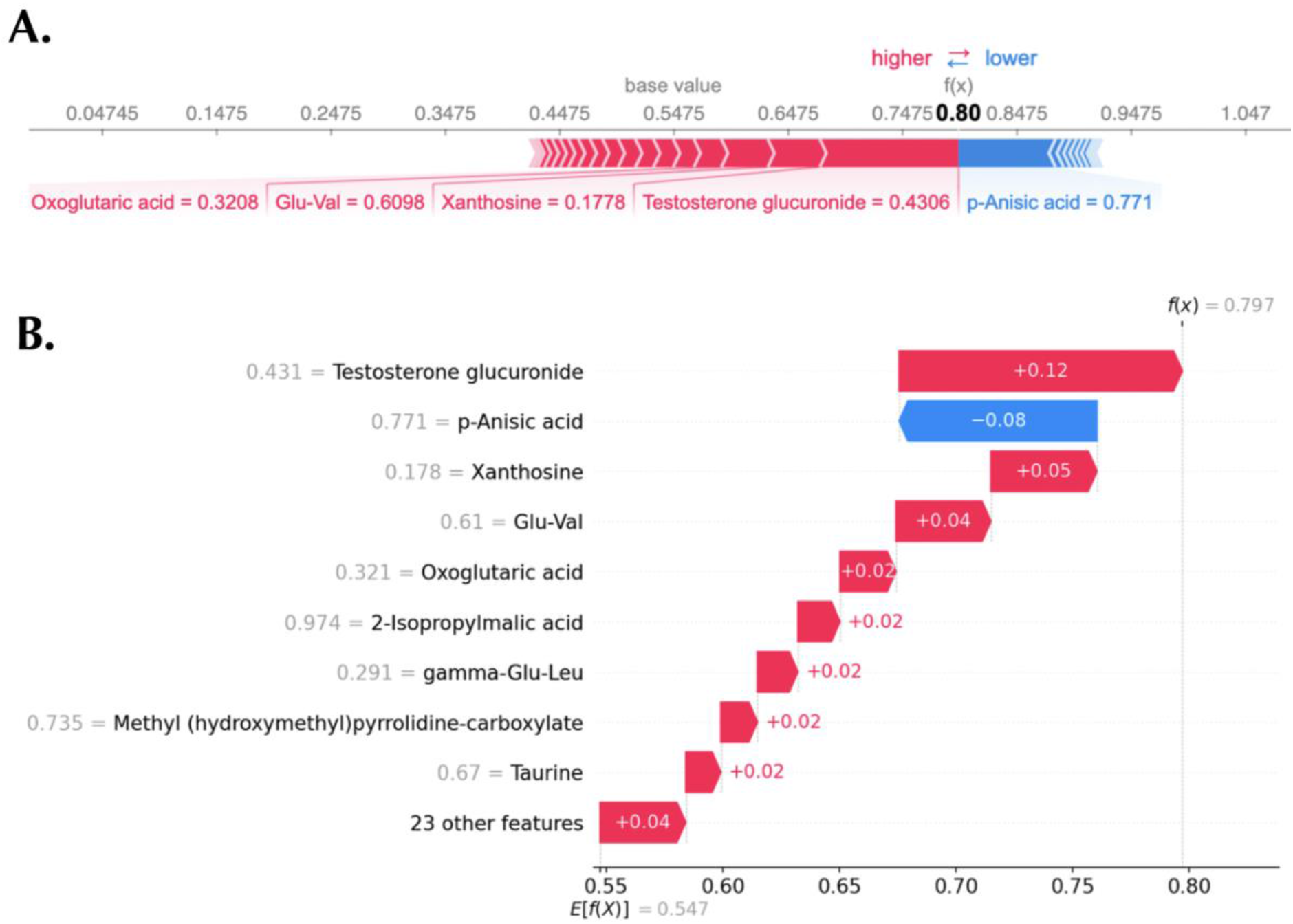
Local explanations of a representative sample. A: Force plot showing a male prediction. B: Waterfall plot displaying the same prediction.

One of the utilities of the SHAP local explanations is the error analysis of individual predictions. The confusion matrix of the test set predictions is shown in Fig 7a, with a true negative rate of 35.48%, a false positive rate of 9.68%, a false negative rate of 6.45%, and a true positive rate of 48.39%. The waterfall plot of two representative true positive samples (male samples successfully classified as males) are shown in Figs 7b and c with a probability output of 0.89 and 0.94, respectively. As in the global explanation, the importance rank of testosterone glucuronide and *p*-anisic acid are conserved. However, a false negative sample (male sample wrongly classified as female) in the test cohort can be seen to be missing testosterone glucuronide in the top 9 important features (Fig 7d). As testosterone glucuronide is a highly ranked metabolomic feature of importance in the true positive samples, this provides a plausible explanation for the model’s wrong classification. In addition, two representative true negative samples (female successfully classified as female) are shown in Figs 7e and f. A low relative abundance of testosterone glucuronide is the most important factor driving the samples’ classification to the female category. Likewise, for the two false-positive samples (females samples wrongly classified as male samples), one of such samples (Fig 7g) has a high relative abundance of testosterone glucuronide, contributing to the wrong classification. On the other hand, another false positive sample (Fig 7h), which has a low relative abundance of the testosterone urinary metabolite, has a probability output of 0.52, just slightly above the female category cut-off value of below 0.5.

**Fig 7.**
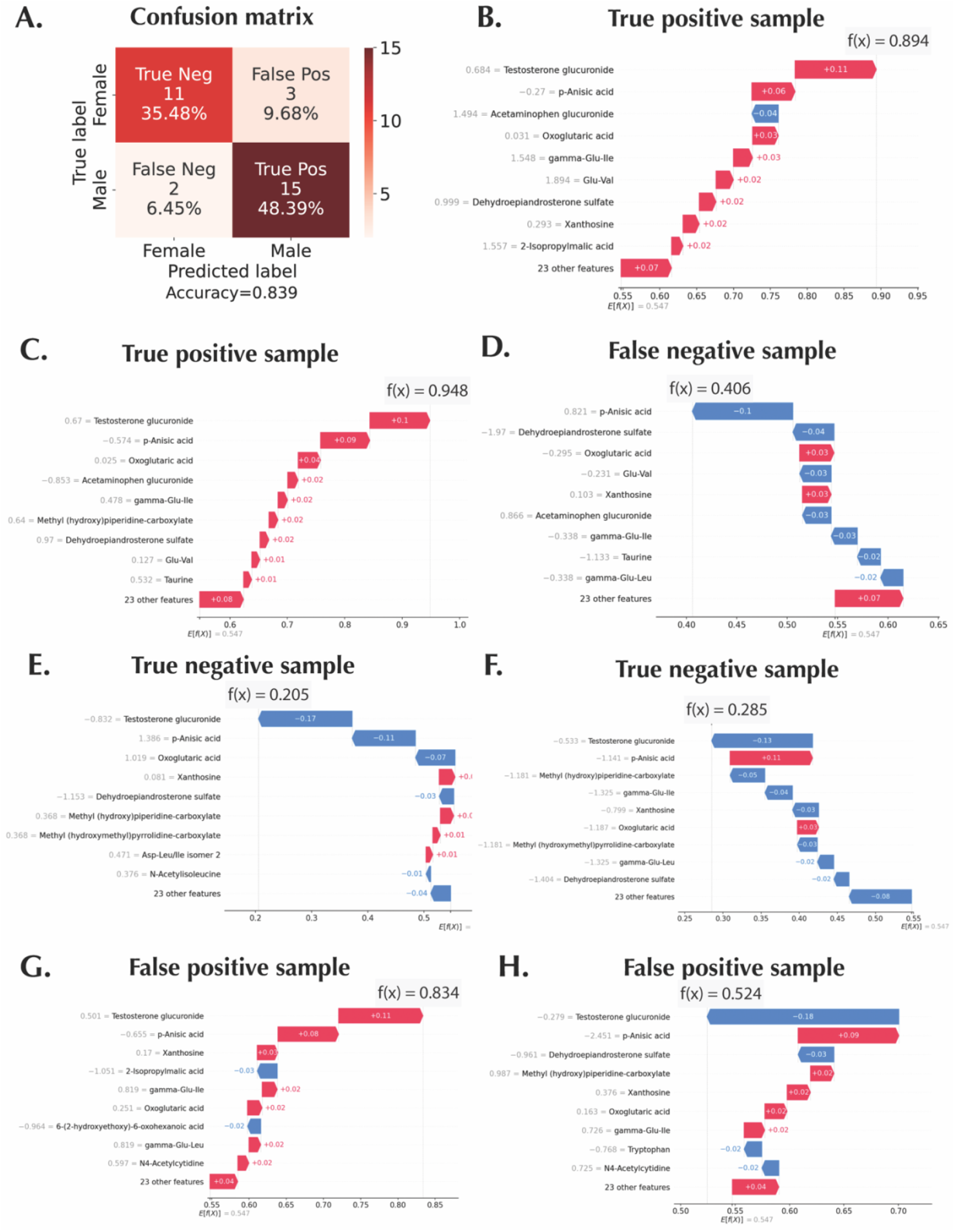
SHAP for error analysis. A: Confusion matrix of the test set for the MTBLS404 dataset. Waterfall plots of, B and C: True positive representative samples. D: A false negative sample. E and F: True negative representative samples. G and H: False positive samples.

## 3 Discussion

In this computational work, after feature selection with fold changes and *T*-test *P* values cut-offs, random forest, gradient boosting, and XGboost were compared with one of the most widely used ML algorithms in the field of metabolomics (PLS-DA) using three publicly available metabolomics datasets. PLS-DA was as predictive as any of the tree-based models on the MTBLS404 dataset, less predictive on at least one of the tree-based models using the MTBLS547. Finally, it was more predictive than all the tree-based models with the ST000369 dataset. Suggesting that multiple ML algorithms should be applied more in metabolomics for classification purposes, especially tree-based models. However, Mendez and co-workers showed that PLS-DA outperformed random forest models in two of the three datasets used in this study, albeit before feature selection and with different sampling methods (33). This might be because random forest models tend to perform poorly when the fraction of relevant features is small in a large variable-sized dataset with a small sample size (34). In this study, relevant features were selected before classification models were built. Furthermore, this paper has shown that there are added benefits to using a tree-based model for classification tasks.

In addition to the compatibility of PLS-DA methods to metabolomics datasets, given that the datasets tend to be linearly separable (33), the interpretability of PLS-DA *via* its VIP score is useful to scientists in the metabolomics field for explaining the model (7). As such, PLS-DA is appealing because it is intrinsically interpretable. However, over the past several years, there had been great stride of achievements in the field of interpretable machine learning, especially in model agnostic methods (35), one of such methods is local interpretable model-agnostic explanations (LIME) (20). The key idea of LIME is that it selects an intrinsically interpretable class of model, and then used that to approximate a black box model locally, therefore interpretations are not necessarily globally faithful. This class of models is called local surrogate models. SHAP unifies LIME and other interpretable machine learning methods with Shapley values to explain both local and global properties leveraging on the unique properties of Shapley values (21). This SHAP computation is called kernel SHAP, and it is computationally expensive and does not calculate exact Shapley values hence the choice of Tree SHAP for the study. On the other hand, Tree SHAP calculates exact Shapley values *via* its conditional expectation method, also it is relatively computationally inexpensive in comparison with kernel SHAP (22).

In this study, the MTBLS404 (sex classification) dataset was used for IML explanations (using a random forest model), because of the combination of the following reasons: 1) the high predictive ROC AUC score of 0.83 on the test set, 2) diversity of errors which allowed for detailed IML utility, and 3) the simplicity of the biological interpretation of the IML results. The VIP score and the mean absolute SHAP values (as shown in the SHAP bar plot) of the MTLBS404 dataset indicate a consensus in the top two most important metabolomic features – testosterone glucuronide and *p*-anisic acid. Apart from that, in general, there are no corresponding matches between both global explanations. However, there is a positive Pearson’s correlation score of 0.50 between the VIP scores and mean absolute SHAP values of the MTLBS404 test set. A perfect correlation score might be expected since the PLS-DA and the random forest models have the same ROC-AUC test set scores. However, it is important to recall that the same predictive performances might not always imply the same explanations, due to the Rashomon effect, which dictates that two models can rely on different features to get the same predictive performance (36). Therefore, consistency in explanations should only be desirable if the models rely on exactly the same relationships to make predictions. On the other hand, the Pearson’s correlation between the SHAP values and random forest’s gini index is 0.99. Since the SHAP analysis was computed using the random forest model, this is an expected result confirming the fidelity of Tree SHAP in computing the feature importance scores.

Going beyond a bar plot of global importance, the summary plot allows for a more detailed explanations of the importance scores using local explanation summaries by showing the impact of the abundance levels of the individual metabolites on the model output. Similar information can also be presented in the SHAP embedding plots, albeit with different visualization, the embedding plots allow the user to see the spread of SHAP values for a particular metabolomic feature. Furthermore, the supervised interpretable hierarchical clustering analysis clustered the model output with explanations similarities. This plot is potentially useful for the discovery of latent population subgroups in a clinical metabolomic study with a rich meta-data (23). However, in this study, the plot allowed for the visualization and hence isolation of testosterone glucuronide as an important marker of sex in urine in this dataset. This is indeed biologically significant as the metabolite is an endogenous, urinary metabolite of testosterone, the principal male sex hormone (37). This is further corroborated *via* the error analysis enabled by the SHAP local interpretation plots as described in detail in the results session.

As shown in this study, Tree SHAP is a desirable post-hoc interpretability method that will enhance the interpretations of metabolomics data analysis, given its local and global interpretations, high expressive power, and high fidelity (14). While SHAP explains predictions, it is always important to note that explanation does not imply causation. This is a general limitation of models and data.

## 4 Methods

### 4.1 Metabolomics Datasets

All the datasets used in this study were deposited on *Metabolomics Workbench* (www.metabolomicsworkbench.org) (38) and *Metabolights* (www.ebi.ac.uk/metabolights) (39, 40) repositories. MTBLS404 and MTBLS547 datasets are from *Metabolomics Workbench,* while ST000369 is from *Metabolights.* Mendez et al 2019 used these datasets in a computational metabolomics study, where the generalized predictive ability of machine learning algorithms was compared (33), and the datasets were converted to the tiny data format in the study for ease of computational analysis. As shown in **Table 1**, biofluids used in the study selected include urine, caecal, and serum. The sample size ranges from 80 to 184, and the metabolic features range from 69 to 184. All machine learning problems are binary classification, and the subject of classification includes sex classification (male *vs.* female) (30), impact of high-fat diet (case *vs.* control) (31), and adenocarcinoma (case *vs.* control) (32).

### 4.2 Machine Learning Models

Partial least squares (PLS) regression models a linear covariance between the feature matrix ***X*** and the response matrix ***Y***. The goal of the algorithm is to predict dependent variables using predictors, and it does so by projecting the data points into a lower dimensional space such that the covariance between response groups are maximized. The projection can be represented mathematically as a PLS coefficient value vector (*B_PLS_*) where predictions are made by 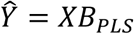. Which is, in essence, a multiple linear regression equation. PLS discriminant analysis (PLS-DA) is the PLS method for binary classification, where 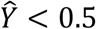 is attributed to a negative classification and 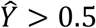 is a positive classification.

Decision trees (DT) are the base learner for tree-based machine learning algorithms. DTs are inverted trees with the root node at the top, the leaves at the bottom, and the internal nodes in between. Root and internal nodes are assigned metabolomic features used for splitting, while the leaves at the end of the tree signify the final predictions. In brief, decision trees split data into branches until the algorithm attains the highest accuracy. This property of DT makes it particularly prone to overfitting, and one of the robust solutions to this problem is the aggregation of decision trees into ensembles such as random forests, gradient boosting, and XGBoost.

Random forest is a bootstrap aggregation (bagging) of decision trees. Each decision trees make its predictions, and the result of the forest of decision trees is made using the majority rules. In the end, bootstrapping increases diversity, while aggregation reduces variance. Despite the strength of random forests, they could be limited by individual trees. If all the trees in the forest make the same mistake, the random forest, in turn, makes that mistake. However, boosting improves on this limitation by making the trees learn from the mistake of the preceding trees; it is this class of tree-based models gradient boosting and extreme gradient boosting (XG-Boost) belongs.

Gradient boosting transforms weak learners into strong ones by iteratively correcting errors, with the base learner being a decision tree. And it does so by fitting the new sequential tree based on the errors of the previous tree. On the other hand, XGBoost is an advanced variant of gradient boosting designed primarily for speed with its speed-enhancing capabilities like approximate and sparsity-aware split-finding, block compression and sharding, parallel computing, and cache-aware computing (41). Also, there can be accuracy gains in using it over gradient boosting and random forests because of the inclusion of regularization in the loss function (41). The regularization parameters help to penalize complexity and prevent overfitting.

### 4.3 Hyperparameter Fine-Tuning

To attempt to improve predictive performance and find a balance between bias and variance, hyper-parameter fine-tuning was carried out (**Table S1-3**). As opposed to parameters learned by the machine learning algorithm, hyper-parameters are set by the user. A linear search for a single hyperparameter or a grid search for two or greater hyperparameters was carried out under 10-fold cross-validation conditions. In PLS-DA, a linear search was conducted for the number of latent variables *(n_components*), the only hyperparameter.

For random forest, hyperparameters considered for tuning include the number of trees in the forest *(n_estimators),* the maximum length of trees *(max_depth),* the maximum number of features to choose from when making a split *(max_features),* the number of samples required before the split can occur (*min_samples_split*), and the minimum number of samples required for a node to be a leaf (*min_samples_leaf*). For XGBoost, hyperparameters considered include maximum depth of tree (*max_depth*), that is the number of branches a tree has; learning rate (*learning_rate*), which limits overfitting by reducing the weight ascribed to each tree to a given percentage; the number of boosted trees in the model (*n_estimators*); *subsample*, which limits the percentage of training samples for each boosting round, and the minimum sum of weight required for a node to split into a child (*min_child_weight*). Finally, for gradient boosting, the following hyperparameters were tuned: *max_depth, subsample, learning_rate, and n_estimators*.

### 4.4 Code and Availability

Programming was carried out using the Python 3.8.12 programming language. SHAP 0.40.0 library was used for SHAP explanations (21, 23, 24). Pandas 1.3.4 was used for data handling (42). Matplotlib 3.5.0 and Seaborn 0.11.2 was used for data visualization (43, 44). Numpy 1.21.1 was used for numerical computation and Sci-kit learn 1.0.2 was used for machine learning. In addition, pyChemometrics 0.1 was used for PLS-DA computations. Jupyter notebook was used as the integrated development environment (IDE). All Jupyter notebooks used in this study can be found here: https://github.com/obifarin/shap-iml-metabolomics

## 5 Acknowledgments

I would like to thank Dr. Arthur S. Edison for feedback about the study. The Georgia Research Alliance supported this computational work.

## 7 Supporting Information

**S1 Fig. Volcano plots for feature selection.** (a), MTBLS404, (b) MTBLS547, (c) ST000369. Significant up (blue) are the metabolomic features that are higher in the males for MTBLS404, high fat diet mice for MTBLS547, and adenocarcinoma lung cancer patients in ST000369, and vice versa.

**S2 Fig. SHAP dependence plots for testosterone glucuronide and γ-glu-leu.** a) Testosterone glucuronide. b) Testosterone glucuronide and γ-glu-leu.

**S1 Table.** Hyperparameters tuned for PLS-DA, random forest, and XGBoost for the train set of the MTBLS404 dataset.

**S2 Table.** Hyperparameters tuned for PLS-DA, random forest, and XGBoost for the train set of the MTBLS547 dataset.

**S3 Table.** Hyperparameters tuned for PLS-DA, random forest, and XGBoost for the train set of the ST000369 dataset.

**Figure S1.**
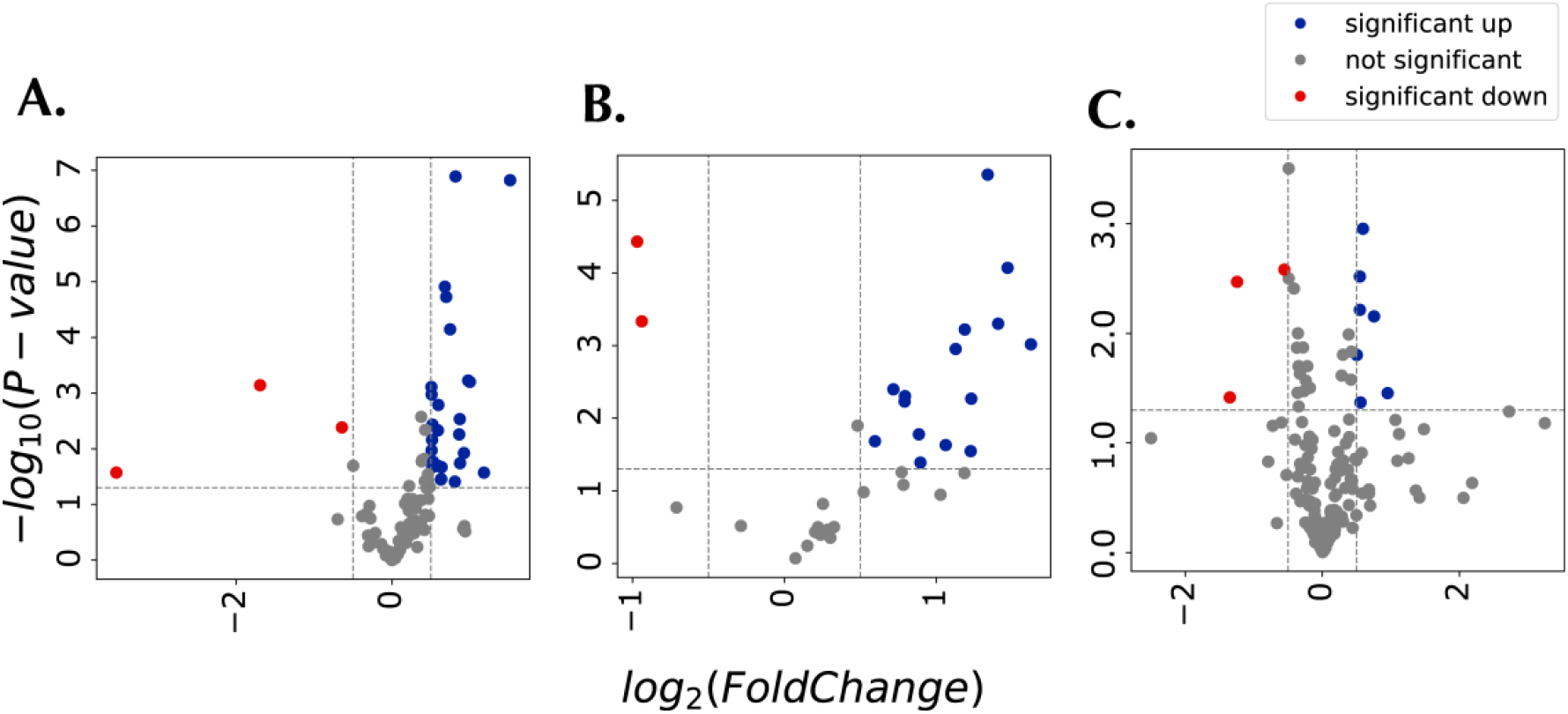
Volcano plots for feature selection. (a), MTBLS404, (b) MTBLS547, (c) ST000369. Significant up (blue) are the metabolomic features that are higher in the males for MTBLS404, high fat diet mice for MTBLS547, and adenocarcinoma lung cancer patients in ST000369, and vice versa.

**Figure S2.**
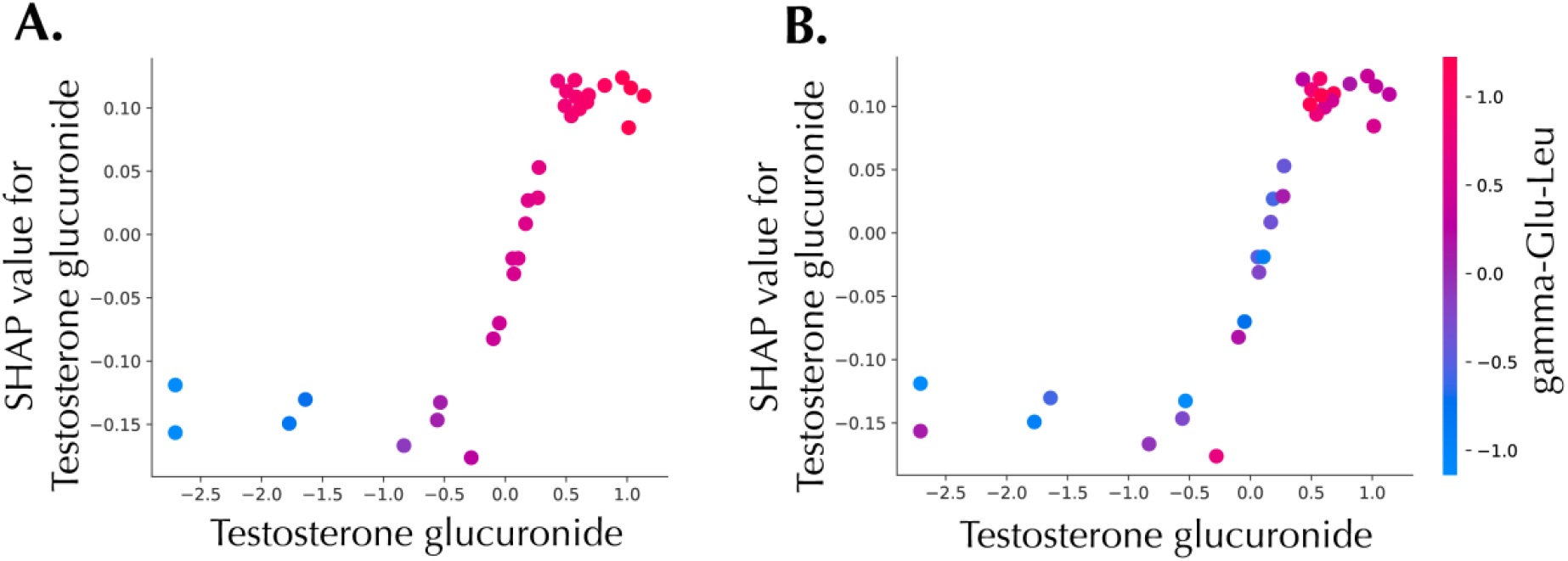
SHAP dependence plots for testosterone glucuronide and γ-glu-leu. a) Testosterone glucuronide. b) Testosterone glucuronide and γ-glu-leu.

**Table S1.**
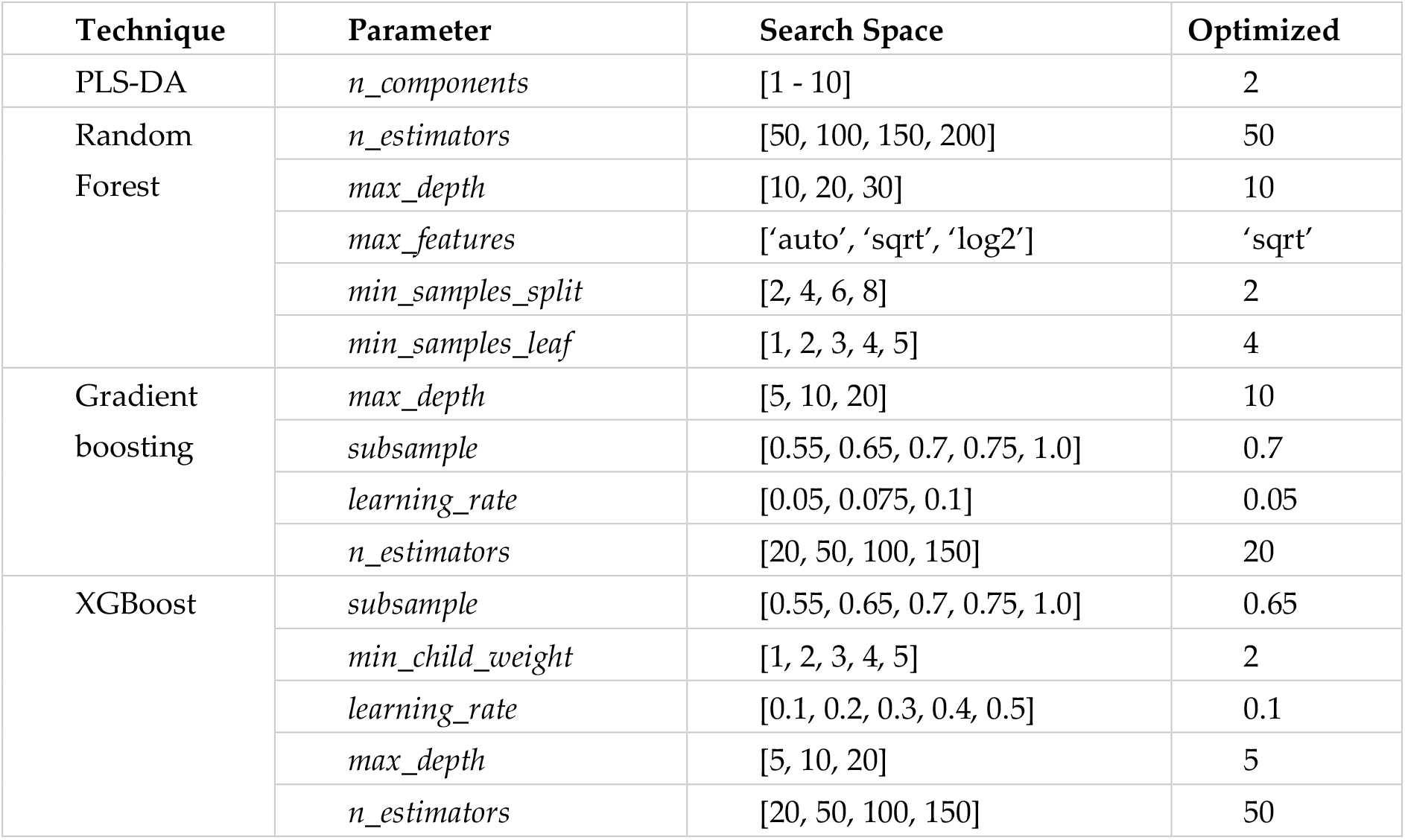
Hyperparameters tuned for PLS-DA, random forest, and XGBoost for the train set of the MTBLS404 dataset.

**Table S2.** Hyperparameters tuned for PLS-DA, random forest, and XGBoost for the train set of the MTBLS547 dataset.

| Technique | Parameter | Search Space | Optimized |
| --- | --- | --- | --- |
| PLS-DA | *n_components* | [1 - 10] | 2 |
| Random Forest | *n_estimators* | [50, 100, 150, 200] | 50 |
|  | *max_depth* | [10, 20, 30] | 10 |
|  | *max_features* | ['auto', 'sqrt', 'log2'] | 'sqrt' |
|  | *min_samples_split* | [2, 4, 6, 8] | 2 |
|  | *min_samples_leaf* | [1, 2, 3, 4, 5] | 1 |
| Gradient boosting | *max_depth* | [5, 10, 20] | 5 |
|  | *subsample* | [0.55, 0.65, 0.7, 0.75, 1.0] | 0.55 |
|  | *learning_rate* | [0.05, 0.075, 0.1] | 0.1 |
|  | *n_estimators* | [20, 50, 100, 150] | 20 |
| XGBoost | *subsample* | [0.5, 0.6, 0.7, 0.8, 0.9, 1] | 0.7 |
|  | *min_child_weight* | [1, 2, 3, 4, 5] | 4 |
|  | *learning_rate* | [0.1, 0.2, 0.3, 0.4, 0.5] | 0.4 |
|  | *max_depth* | [1, 2, 3, 4, 5, None] | 2 |
|  | *n_estimators* | [2, 25, 50, 75, 100] | 50 |

**Table S3.** Hyperparameters tuned for PLS-DA, random forest, and XGBoost for the train set of the ST000369 dataset.

| Technique | Parameter | Search Space | Optimized |
| --- | --- | --- | --- |
| PLS-DA | *n_components* | [1 - 10] | 1 |
| Random Forest | *n_estimators* | [50, 100, 150, 200] | 150 |
|  | *max_depth* | [2, 5, 10] | 5 |
|  | *max_features* | ['auto', 'sqrt', 'log2'] | 'sqrt' |
|  | *min_samples_split* | [2, 4, 6, 8] | 4 |
|  | *min_samples_leaf* | [1, 2, 3, 4, 5] | 1 |
| Gradient boosting | *max_depth* | [5, 10, 20, 25] | 20 |
|  | *subsample* | [0.55, 0.65, 0.7, 0.75, 1.0] | 0.65 |
|  | *learning_rate* | [0.05, 0.075, 0.1] | 0.075 |
|  | *n_estimators* | [20, 50, 100, 150] | 50 |
| XGBoost | *subsample* | [0.5, 0.6, 0.7, 0.8, 0.9, 1] | 0.8 |
|  | *min_child_weight* | [1, 2, 3, 4, 5] | 1 |
|  | *learning_rate* | [0.1, 0.2, 0.3, 0.4, 0.5] | 0.3 |
|  | *max_depth* | [1, 2, 3, 4, 5, None] | 1 |
|  | *n_estimators* | [2, 25, 50, 75, 100] | 50 |

## References

1. Nicholson JK, Lindon JC. Systems biology: Metabonomics. Nature. 2008;455(7216):1054–6.

2. Bzdok D, Altman N, Krzywinski M. Statistics versus machine learning. Nat Methods. 2018;15(4):233–4.

3. Bzdok D, Krzywinski M, Altman N. Points of Significance: Machine learning: a primer. Nat Methods. 2017;14(12):1119–20.

4. Smolinska A, Blanchet L, Buydens LM, Wijmenga SS. NMR and pattern recognition methods in metabolomics: from data acquisition to biomarker discovery: a review. Anal Chim Acta. 2012;750:82–97.

5. Grissa D, Petera M, Brandolini M, Napoli A, Comte B, Pujos-Guillot E. Feature Selection Methods for Early Predictive Biomarker Discovery Using Untargeted Metabolomic Data. Front Mol Biosci. 2016;3:30.

6. Worley B, Powers R. Multivariate Analysis in Metabolomics. Curr Metabolomics. 2013;1(1):92–107.

7. Gromski PS, Muhamadali H, Ellis DI, Xu Y, Correa E, Turner ML, et al. A tutorial review: Metabolomics and partial least squares-discriminant analysis--a marriage of convenience or a shotgun wedding. Anal Chim Acta. 2015;879:10–23.

8. Ruiz-Perez D, Guan H, Madhivanan P, Mathee K, Narasimhan G. So you think you can PLS-DA? BMC Bioinformatics. 2020;21(Suppl 1):2.

9. Galindo-Prieto B, Eriksson L, Trygg J. Variable influence on projection (VIP) for orthogonal projections to latent structures (OPLS). J Chemom. 2014;28:623–32.

10. Wu L, Huang R, Tetko IV, Xia Z, Xu J, Tong W. Trade-off Predictivity and Explainability for Machine-Learning Powered Predictive Toxicology: An in-Depth Investigation with Tox21 Data Sets. Chem Res Toxicol. 2021;34(2):541–9.

11. London AJ. Artificial Intelligence and Black-Box Medical Decisions: Accuracy versus Explainability. Hastings Cent Rep. 2019;49(1):15–21.

12. Mosconi F, Julou T, Desprat N, Sinha DK, Allemand J-F, Croquette V, et al. Some nonlinear challenges in biology. Nonlinearity. 2008;21(8):131–47.

13. Ribeiro MT, Singh S, Guestrin C. Model-Agnostic Interpretability of Machine Learning. arXiv. 2016:arXiv:1606.05386.

14. Molnar C. Interpretable machine learning. A Guide for Making Black Box Models Explainable. 2019.

15. Friedman JH. Greedy function approximation: A gradient boosting machine. Ann Statist. 2001;29(5):1189–232.

16. Goldstein A, Kapelner A, Bleich J, Pitkin E. Peeking Inside the Black Box: Visualizing Statistical Learning With Plots of Individual Conditional Expectation. J Comput Graph Stat. 2015;24(1):44–65.

17. Apley DW, Zhu J. Visualizing the Effects of Predictor Variables in Black Box Supervised Learning Models. arXiv. 2019:arXiv:1612.08468.

18. Breiman L. Random Forests. Machine Learning. 2001;45:5–32.

19. Fisher A, Rudin C, Dominici F. All Models are Wrong, but Many are Useful: Learning a Variable’s Importance by Studying an Entire Class of Prediction Models Simultaneously. arXiv. 2019:arXiv:1801.01489v5.

20. Ribeiro MT, Singh S, Guestrin C. “Why Should I Trust You?”: Explaining the Predictions of Any Classifier. arXiv. 2016:arXiv:1602.04938v3.

21. Lundberg SM, Lee SI. A Unified Approach to Interpreting Model Predictions. arXiv. 2017:arXiv:1705.07874v2.

22. Lundberg SM, Erion G, Lee SI. Consistent Individualized Feature Attribution for Tree Ensembles. arXiv. 2018:arXiv:1802.03888v3.

23. Lundberg SM, Erion G, Chen H, DeGrave A, Prutkin JM, Nair B, et al. From Local Explanations to Global Understanding with Explainable AI for Trees. Nat Mach Intell. 2020;2(1):56–67.

24. Lundberg SM, Nair B, Vavilala MS, Horibe M, Eisses MJ, Adams T, et al. Explainable machine-learning predictions for the prevention of hypoxaemia during surgery. Nat Biomed Eng. 2018;2(10):749–60.

25. Rodriguez-Perez R, Bajorath J. Interpretation of machine learning models using shapley values: application to compound potency and multi-target activity predictions. J Comput Aided Mol Des. 2020;34(10):1013–26.

26. Rodriguez-Perez R, Bajorath J. Interpretation of Compound Activity Predictions from Complex Machine Learning Models Using Local Approximations and Shapley Values. J Med Chem. 2020;63(16):8761–77.

27. Cha Y, Shin J, Go B, Lee DS, Kim Y, Kim T, et al. An interpretable machine learning method for supporting ecosystem management: Application to species distribution models of freshwater macroinvertebrates. J Environ Manage. 2021;291:112719.

28. Liu X, Locasale JW. Metabolomics: A Primer. Trends Biochem Sci. 2017;42(4):274–84.

29. Shapley LS. A value for n-person games. Contributions to the Theory of Games. 1953;2(28):307–17.

30. Thevenot EA, Roux A, Xu Y, Ezan E, Junot C. Analysis of the Human Adult Urinary Metabolome Variations with Age, Body Mass Index, and Gender by Implementing a Comprehensive Workflow for Univariate and OPLS Statistical Analyses. J Proteome Res. 2015;14(8):3322–35.

31. Zheng X, Huang F, Zhao A, Lei S, Zhang Y, Xie G, et al. Bile acid is a significant host factor shaping the gut microbiome of diet-induced obese mice. BMC Biol. 2017;15(1).

32. Fahrmann JF, Kim K, DeFelice BC, Taylor SL, Gandara DR, Yoneda KY, et al. Investigation of metabolomic blood biomarkers for detection of adenocarcinoma lung cancer. Cancer Epidemiol Biomarkers Prev. 2015;24(11):1716–23.

33. Mendez KM, Reinke SN, Broadhurst DI. A comparative evaluation of the generalised predictive ability of eight machine learning algorithms across ten clinical metabolomics data sets for binary classification. Metabolomics. 2019;15(12):150.

34. Hastie T, Tibshirani R, Friedman J. The elements of statistical learning (2nd ed.): New York: Springer; 2009.

35. Linardatos P, Papastefanopoulos V, Kotsiantis S. Explainable AI: A Review of Machine Learning Interpretability Methods. Entropy (Basel). 2020;23(1).

36. Breiman L. Statistical Modeling: The Two Cultures (with comments and a rejoinder by the author). Statist Sci. 2001;16(3):199–231.

37. Zumoff B, Rosenfeld RS, Friedman M, Byers SO, Rosenman RH, Hellman L. Elevated daytime urinary excretion of testosterone glucuronide in men with the type A behavior pattern. Psychosom Med. 1984;46(3):223–5.

38. Sud M, Fahy E, Cotter D, Azam K, Vadivelu I, Burant C, et al. Metabolomics Workbench: An international repository for metabolomics data and metadata, metabolite standards, protocols, tutorials and training, and analysis tools. Nucleic Acids Res. 2016;44(D1):D463–70.

39. Haug K, Salek RM, Conesa P, Hastings J, de Matos P, Rijnbeek M, et al. MetaboLights--an open-access general-purpose repository for metabolomics studies and associated meta-data. Nucleic Acids Res. 2013;41(Database issue):D781–6.

40. Haug K, Cochrane K, Nainala VC, Williams M, Chang J, Jayaseelan KV, et al. MetaboLights: a resource evolving in response to the needs of its scientific community. Nucleic Acids Res. 2020;48(D1):D440–D4.

41. Chen T, Guestrin C. XGBoost: A Scalable Tree Boosting System. Proceedings of the 22nd ACM SIGKDD International Conference on Knowledge Discovery and Data Mining. 2016.

42. McKinney W. Data structures for statistical computing in python. Proceedings of the 9th Python in Science Conference. 2010;445:56–61.

43. Hunter JD. Matplotlib: A 2D graphics environment. Computing in Science & Engineering. 2007;9(3):90–5.

44. Waskom ML. Seaborn: statistical data visualization. Journal of Open Source Software. 2021;6(60).

